# Coordinated Membrane Deformation Driven by a Minimal Set of *Spiroplasma* MreB Isoforms

**DOI:** 10.64898/2026.04.13.718137

**Authors:** Takahiro Mitani, Taiki Nishimura, Hana Kiyama, Ahsan Ali, Masahito Hayashi, Kingo Takiguchi, Makoto Miyata, Ikuko Fujiwara

## Abstract

*Spiroplasma* are wall-less helical bacteria that swim by switching the handedness of their helices. In most bacteria, MreB, a bacterial actin, forms relatively static filaments beneath the membrane to organize cell wall synthesis. In contrast, *Spiroplasma eriocheiris* encodes five MreB isoforms (SpeMreBs), and swimming requires a pair of isoforms, SpeMreB5 together with SpeMreB4 (or the related isoform SpeMreB1) (1) Yet how these MreBs generate force and membrane deformation remains unclear. To examine the membrane-deforming activities of SpeMreBs, we demonstrated a simple reconstitution system using the non-motile synthetic bacterium JCVI-syn3B and purified SpeMreBs expressed in *Escherichia coli*. Lysates of syn3B expressing SpeMreB5 deformed liposomes in a concentration-dependent manner. In contrast, lysates co-expressing both SpeMreB5 and SpeMreB4 showed a plateau in the frequency of deformation, suggesting that SpeMreB4 suppresses membrane deformation driven by SpeMreB5. Deformed liposomes exhibited either fluctuating or stable behaviors. ATP depletion changed both the frequency and behavior of deformation, indicating that membrane remodeling depends on the nucleotide state of SpeMreBs. Reconstitution with purified SpeMreBs from *E. coli* confirmed that SpeMreB5 alone deforms membranes, whereas SpeMreB1, a member of the same class as SpeMreB4, suppresses deformation. These results suggest that membrane shape in *Spiroplasma* is dynamically regulated by antagonistic interactions among isoforms of SpeMreBs isoforms and their nucleotide-dependent assembly states.

## Introduction

*Spiroplasma* are thin helical bacteria belonging to the class Mollicutes. *Spiroplasma* infect insects, crustaceans, and plants (2–4). Phylogenetic analyses have recently revised the taxonomy of Mollicutes, including Mycoplasma and related genera, placing *Spiroplasma* as a distinct lineage within wall-less bacteria (5). Unlike most motile bacteria use flagella for moving, *Spiroplasma* lack flagella and instead move by switching the handedness of their helical cell bodies (6, 7). This motility arises from a localized change in helicity, termed a kink, that forms near one end of the cell and travels along the cell axis (7, 8). As the kink moves along the cell axis, the helical segments on either side rotate in opposite directions, generating opposing torques. These coordinated rotations propel the cell forward. Mechanical models based on helical rotation and torque balance have been proposed to explain this propulsion mechanism (9, 10). The waiting time between kink events may depend on intracellular ATP levels (11). However, the mechanisms underlying *Spiroplasma* motility remain unclear.

Without peptidoglycan cell wall that maintains cell shape in most bacteria (12, 13), *Spiroplasma* possess a ribbon-like cytoskeletal structure beneath the membrane that runs along the cell axis and consists of fibril proteins and MreB filaments (14, 15). MreB is a bacterial actin that forms antiparallel double protofilament structures distinct from eukaryotic actin filaments. Structural studies indicate that curvature and spatial arrangement of MreB filaments can generate mechanical stresses on membranes (16). Most bacteria encode a single MreB protein, but *Spiroplasma* possess five MreB isoforms (SMreB1–5) that are grouped into three classes: SMreB1/SMreB4, SMreB2/SMreB5, and SMreB3 (1, 17). In this study, the isoforms from *Spiroplasma eriocheiris* are designated SpeMreB1–5.

A minimal synthetic bacterium, JCVI-syn3B (syn3B), derived from *Mycoplasma mycoides*, exhibits a simple spherical and non-motile phenotype (18). Expression of each isoform of SpeMreB in the syn3B revealed that individual SpeMreBs do not significantly alter cell morphology. However, co-expression of one isoform from the SpeMreB1/MreB4 class together with one from the SpeMreB2/MreB5 class produces elongated helical cells capable of swimming (1).

These results suggest that a minimal set of SpeMreB isoforms is sufficient to reconstitute *Spiroplasma*-like helical motility in cells. This raises a fundamental question: can such minimal components directly generate membrane deformation and motility in a simplified cell-like system, or are additional cellular factors required? Addressing this requires a bottom-up approach to determine whether SpeMreBs alone are sufficient to deform membranes and produce motile behavior in a reconstituted system.

Biochemical studies using purified SpeMreBs expressed in Escherichia coli further show that the isoforms exhibit distinct nucleotide-dependent polymerization properties and interactions (19, 20). Together with observations of kink propagation, these findings suggest that the *Spiroplasma* cytoskeleton is not static but rather forms a dynamic ATP-dependent system with functional specialization among isoforms (6, 21). However, the molecular mechanism by which SpeMreB generates helical shape and drives helical switching remains largely unknown (6, 22, 23).

To address these questions, we used a simple reconstitution approach with liposomes. Previous studies have shown that alignment of actin filaments within liposomes can deform membranes from spherical to elongated shapes (24), and that liposomes containing actin and myosin exhibit contractile behaviors associated with cytoskeletal reorganization (25). Using lysates from syn3B cells expressing SpeMreBs, we analyzed ATP-dependent membrane deformation in liposomes. We then reconstituted liposomes with purified SpeMreBs produced in *E. coli* to dissect the contributions of individual isoforms. Although the reconstituted liposomes did not exhibit directional motility, they displayed distinct deformation behaviors. These experiments revealed that SpeMreB5 actively deforms the membrane, whereas SpeMreB4/MreB1-class isoforms modulate the patterns and dynamics of membrane deformation. Moreover, these activities depend on the nucleotide state of the SpeMreBs, suggesting that cooperative interactions among SpeMreBs can drive ATP-dependent membrane remodeling.

## Result

### Liposomes encapsulating syn3B lysates expressing *Spiroplasma* MreB deform their shape

To test whether *Spiroplasma* MreB can induce membrane deformation, we encapsulated lysates of the minimal synthetic bacterium, syn3B into cell-sized giant liposomes made from dioleoyl-*sn*-glycero-3-phosphatidylcholine (DOPC liposomes) using a water-in-oil (W/O) emulsion method (24). Throughout this study, lysates prepared from syn3B cells expressing SpeMreBs are referred to as SpeMreB5^syn3B lysate^ or SpeMreB5/SpeMreB4^syn3B lysate^. Co-expression of SpeMreB4 and SpeMreB5 is known to be required for helical cell shape and swimming motility in Spiroplasma (1).

Liposome shape was monitored over time by phase-contrast and fluorescence microscopy. As a control, liposomes encapsulating lysates from syn3B cells with no SpeMreB expression were examined. Most control liposomes remained spherical. Fluorescent dye added to the lysate confirmed efficient encapsulation of syn3B cytoplasmic contents inside the liposomes **(Fig. 1A, Movie S1 and S2)**. In contrast, liposomes encapsulating SpeMreB5-mCherry^syn3B lysate^, or both SpeMreB5-mCherry^syn3B lysate^ and SpeMreB4^syn3B lysate^, frequently exhibited non-spherical shapes **(Fig. 1B, 1C, Movie S3-S6)**. Incorporation of SpeMreB5^syn3B lysate^ into liposomes was verified by mCherry fluorescence. To quantify the frequency and extent of deformation, the long and short axes of individual liposomes were measured, and their aspect ratios were plotted **(Fig. 1D–1F)**. Liposomes located along the diagonal have nearly equal axes and therefore remain spherical. In this analysis, liposomes with an aspect ratio ≥ 0.95 were classified as spherical, whereas those with an aspect ratio < 0.95 were considered deformed. Plots of aspect ratio at different lysate concentrations **(Fig. S1)** were used to calculate the fraction of deformed liposomes **(Fig. 1G)**. In control lysates lacking SpeMreB expression, the fraction of deformed liposomes varied between ∼10% and 30% and did not show a clear dependence on lysate concentration. In contrast, liposomes containing SpeMreB5^syn3B lysate^ showed a linear increase in deformation frequency with increasing lysate concentration, reaching ∼71% of liposomes. These observations indicate that SpeMreB5^syn3B lysate^ promotes membrane deformation. Notably, when SpeMreB4^syn3B lysate^ was present together with SpeMreB5^syn3B lysate^, the fraction of deformed liposomes increased with concentration but plateaued at ∼60% above OD_600_ ≈ 0.2. This plateau suggests that SpeMreB4^syn3B lysate^ limits or modulates membrane deformation driven by SpeMreB5^syn3B lysate^.

**Figure 1.**
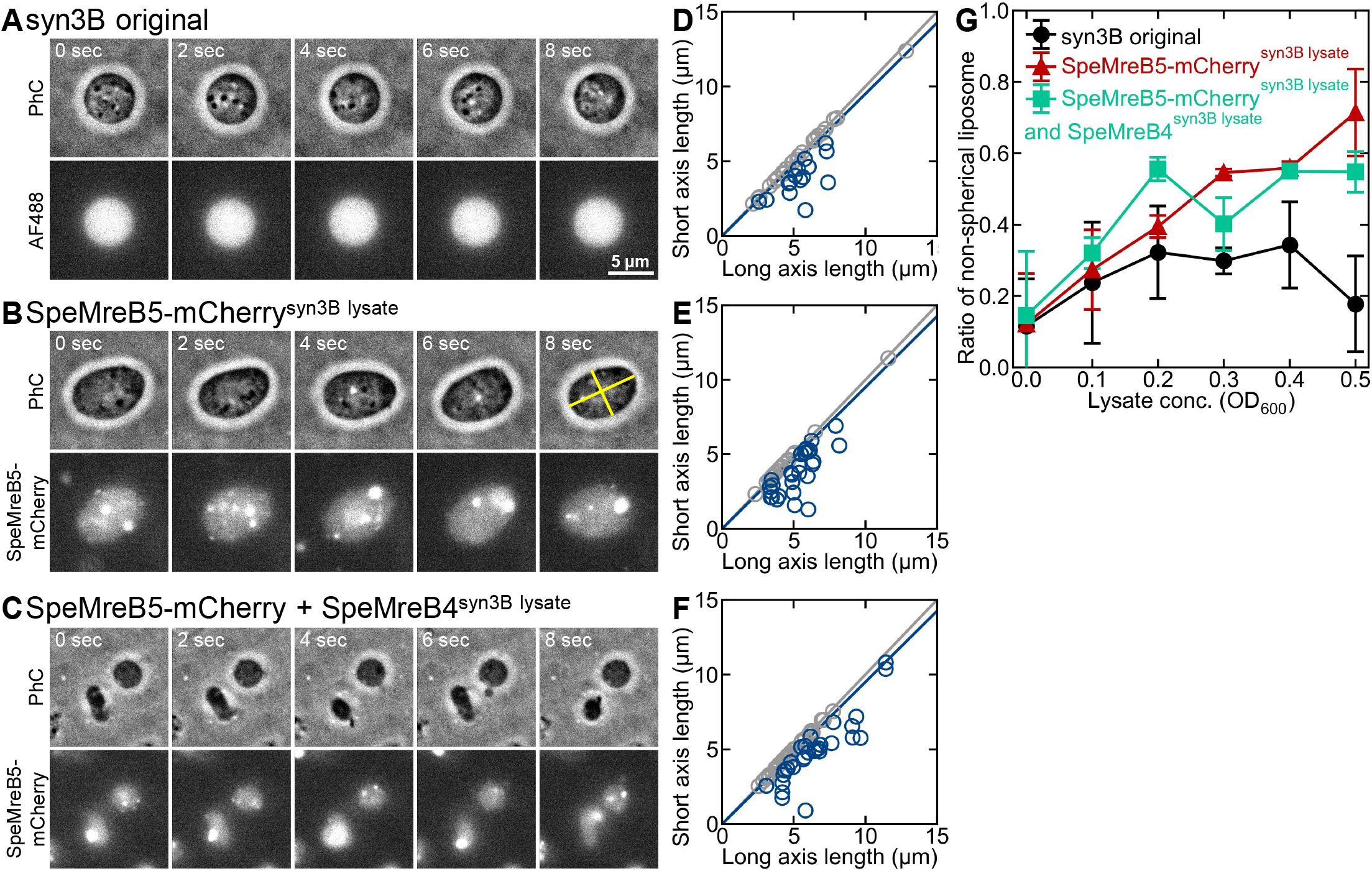
syn3B lysates containing *Spiroplasma* MreB actively deform liposomes. Liposomes encapsulating syn3B lysates expressing SpeMreB proteins change shape over time. (A–C) Time-lapse images of liposomes encapsulating different syn3B lysates **(see Movies S1– S3**). (A) Control lysate from syn3B cells with no SpeMreB expression. (B) Lysate from syn3B cells expressing SpeMreB5-mCherry. (C) Lysate from syn3B cells co-expressing SpeMreB5-mCherry and SpeMreB4. All lysate concentrations were adjusted to OD_600_ = 0.3 before encapsulation. The upper panels show phase-contrast images and the lower panels show fluorescence images of Alexa Fluor 488 (A) or SpeMreB5-mCherry (B,C). Time is indicated in the phase-contrast images (images were not acquired simultaneously). The scale bar of 5 μm is shown in panel A and applies to all panels. The yellow lines in the phase-contrast image in (B) illustrate representative measurements of the long and short axes. (D–F) Scatter plots of long-axis versus short-axis lengths for individual liposomes under each condition. The black diagonal indicates an aspect ratio of 1 (perfect sphere), and the blue line indicates an aspect ratio of 0.95. Liposomes with an aspect ratio <0.95 are shown in blue and were classified as deformed. (G) Fraction of deformed liposomes as a function of lysate concentration. Data represents two independent experiments, each analyzing more than 50 liposomes. Control lysate lacking SpeMreB expression (black) showed little dependence on concentration. Lysates expressing SpeMreB5-mCherry (red) produced a concentration-dependent increase in deformation. Lysates co-expressing SpeMreB5-mCherry and SpeMreB4 (green) reached a plateau above OD_600_ ≈ 0.2. (see **Mov. S1-S3)**.

### Nucleotide state controls the frequency of liposome deformation and membrane dynamics

Membrane deformation driven by SpeMreBs^syn3B lysate^ is expected to depend on the nucleotide state of the proteins. Previous sedimentation assays using purified proteins expressed in *E. coli* showed that SpeMreB1, which belongs to the same class as SpeMreB4, increases its own polymerized structure in the ADP-bound state, while reducing it in the presence of SpeMreB5 (20, 26). Based on this observation, we tested whether nucleotide depletion alters membrane deformation in liposomes. We used syn3B lysates expressing both SpeMreB5–mCherry and SpeMreB4. ATP in syn3B lysates was depleted by the addition of hexokinase with glucose. MgCl_2_ present in the buffer was sufficient to support hexokinase activity, and the resulting liposome behavior and dynamics were analyzed. Two distinct dynamic behaviors of deformed liposomes were observed during 10 s observations: liposomes that maintained a stable deformed shape and liposomes whose membranes fluctuated while remaining deformed **(Fig. 2A; Movies S7 and S8)**. The fraction of liposomes with stable deformation increased with lysate concentration but decreased when the concentration exceeded OD_600_ ≈ 0.2 **(Fig. 2B)**. This concentration dependence was similar with or without hexokinase, although ATP depletion increased the fraction of these liposomes from ∼20% to ∼30%. In contrast, liposomes showing membrane fluctuations increased in frequency with increasing lysate concentration and reached a plateau above OD_600_ ≈ 0.3. This tendency was also observed both before and after ATP depletion of the lysate, although the fraction increased slightly after ATP depletion.

**Figure 2.**
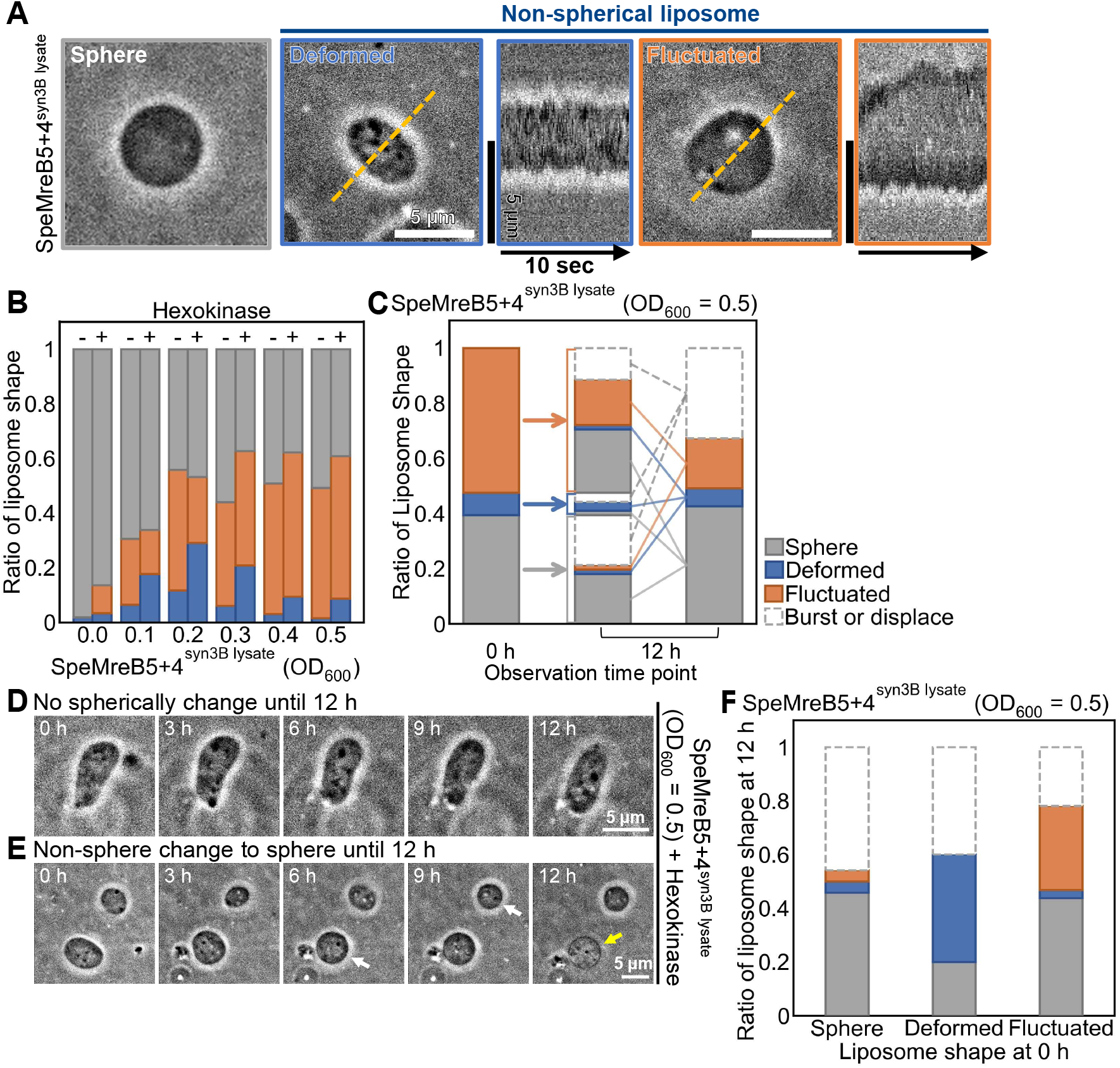
ATP depletion alters the pattern of liposome deformation and membrane dynamics driven by SpeMreBs ^syn3B lysate^. (A) Representative patterns of liposome deformation encapsulating syn3B lysates expressing SpeMreB5-mCherry and SpeMreB4. Liposomes were categorized as spherical (gray), stably deformed without membrane fluctuations (blue), or deformed with membrane fluctuations (orange). Kymographs generated along the yellow dashed lines are shown on the right (**Movies S7 and S8**). Scale bar, 5 μm. (B) Frequency of each liposome pattern as a function of lysate concentration under conditions with (+) or without (-) hexokinase with glucose. Spherical liposomes (gray) dominated at low lysate concentrations. Stably deformed liposomes (blue) increased around OD_600_ = 0.2–0.3. Liposomes showing membrane fluctuations (orange) increased with lysate concentration and plateaued above OD_600_ ≈ 0.3. Similar trends were observed before and after ATP depletion. (C) Distribution of liposome patterns immediately after hexokinase addition (0 h, left column) and after 12 h incubation (center and right columns) at OD_600_ = 0.5. The central column and arrows indicate the pattern of change based on liposome behavior at 0 h. The right column and lines indicate the distribution of the overall liposome patterns after 12 h. Although some liposomes ruptured during the incubation, the overall patterns distribution remained broadly similar. Colors correspond to those in (B). (D,E) Representative examples of liposomes tracked for 12 h after hexokinase addition (**Movie S9**). Liposomes maintaining stable deformation (D) and liposomes transitioning from deformed to spherical shapes (E, white arrow) are shown. Liposomes that lost internal contents due to rupture are also indicated (yellow arrow). (F) Transition analysis of liposome patterns over 12 h following hexokinase addition. Histograms indicate the initial pattern of each liposome. Some liposomes disappeared during the observation period due to rupture (dashed boxes). Approximately 40% of liposomes initially exhibiting membrane fluctuations transitioned to spherical shapes, suggesting that sustained deformation and membrane fluctuations depend on ATP. Colors correspond to those in (B,C) (**Movie S10**).

To examine longer-term effects of ATP depletion, liposome behavior was monitored for 12 h after hexokinase addition **(Fig. 2C)**. Comparison of the distributions at 0 h and 12 h **(Fig. 2C, left and right)** shows that a substantial fraction of liposomes retained their initial shapes. Some liposomes, however, underwent changes in behavior, including transitions from fluctuating to spherical states **(Fig. 2C, center)**. We first considered changes in the overall distribution. The total number of liposomes decreased by ∼30% during this period, largely due to liposome rupture **(Fig. 2C, right)**. The fraction of spherical liposomes remained approximately constant at ∼40% **(Fig. 2C, gray)**, and the fraction of stably deformed liposomes remained ∼7% **(Fig. 2C, blue)**. In contrast, the fraction of fluctuating liposomes decreased substantially from ∼50% to ∼20% **(Fig. 2C, orange)**. Next, we examined how individual liposomes contributed to these changes by tracking the same liposomes over time **(Fig. 2C center, 2D, 2E; Movies S9 and S10)**. Transition analysis revealed that among liposomes initially exhibiting stable deformation, ∼40% remained unchanged, ∼40% disappeared, and ∼20% became spherical **(Fig. 2C center and Fig. 2F center, blue)**. Among liposomes initially exhibiting membrane fluctuations, ∼45% became spherical over 12 h, whereas ∼30% retained fluctuations and ∼20% disappeared **(Fig. 2C center and 2F right, orange)**. A small fraction (∼5%) transitioned to stable deformation without fluctuations **(Fig. 2F right, blue)**. These observations indicate that ATP depletion alters the stability of membrane deformation induced by SpeMreBs. In particular, fluctuating liposomes frequently reverted to spherical shapes after ATP depletion, suggesting that ATP-dependent interactions among SpeMreBs contribute to sustained membrane deformation and membrane fluctuations. One possible explanation based on a previous paper (20) is that, in the ADP-bound state, enhanced polymerized form of SpeMreB1-class proteins destabilizes SpeMreB5 filaments, weakening the forces that maintain dynamic membrane deformation. In contrast, the persistence of some stably deformed liposomes suggests that ATP-independent interactions between SpeMreBs and the membrane can also maintain certain deformed states.

### Purified SpeMreBs exert opposing effects on liposome deformation

To determine whether SpeMreBs directly contribute to liposome deformation, we examined the effects of purified proteins expressed in *E. coli* (SpeMreBs^purified^). Because SpeMreB4^purified^ is difficult to purify (20), we used SpeMreB1^purified^ as a representative of the same functional class. Previous work showed that SpeMreB1^purified^ polymerizes in an ATP-dependent manner and interacts with SpeMreB5^purified^ (20). For these experiments, SpeMreB1 was expressed as an N-terminal fusion with the solubility tag Protein S from *Myxococcus xanthus* (PrS-SpeMreB1^purified^), which was retained for solubility (27). PrS-SpeMreBs^purified^ were encapsulated in DOPC liposomes using the W/O emulsion transfer method **(Fig. 3A–D and Movie S11–S15)**. Under all conditions tested, the frequency of deformed liposomes was lower than that observed with SpeMreBs^syn3B lysate^ **(Fig. 3E–3L)**, suggesting that lipids or other cellular components in the lysate facilitate membrane deformation.

**Figure 3.**
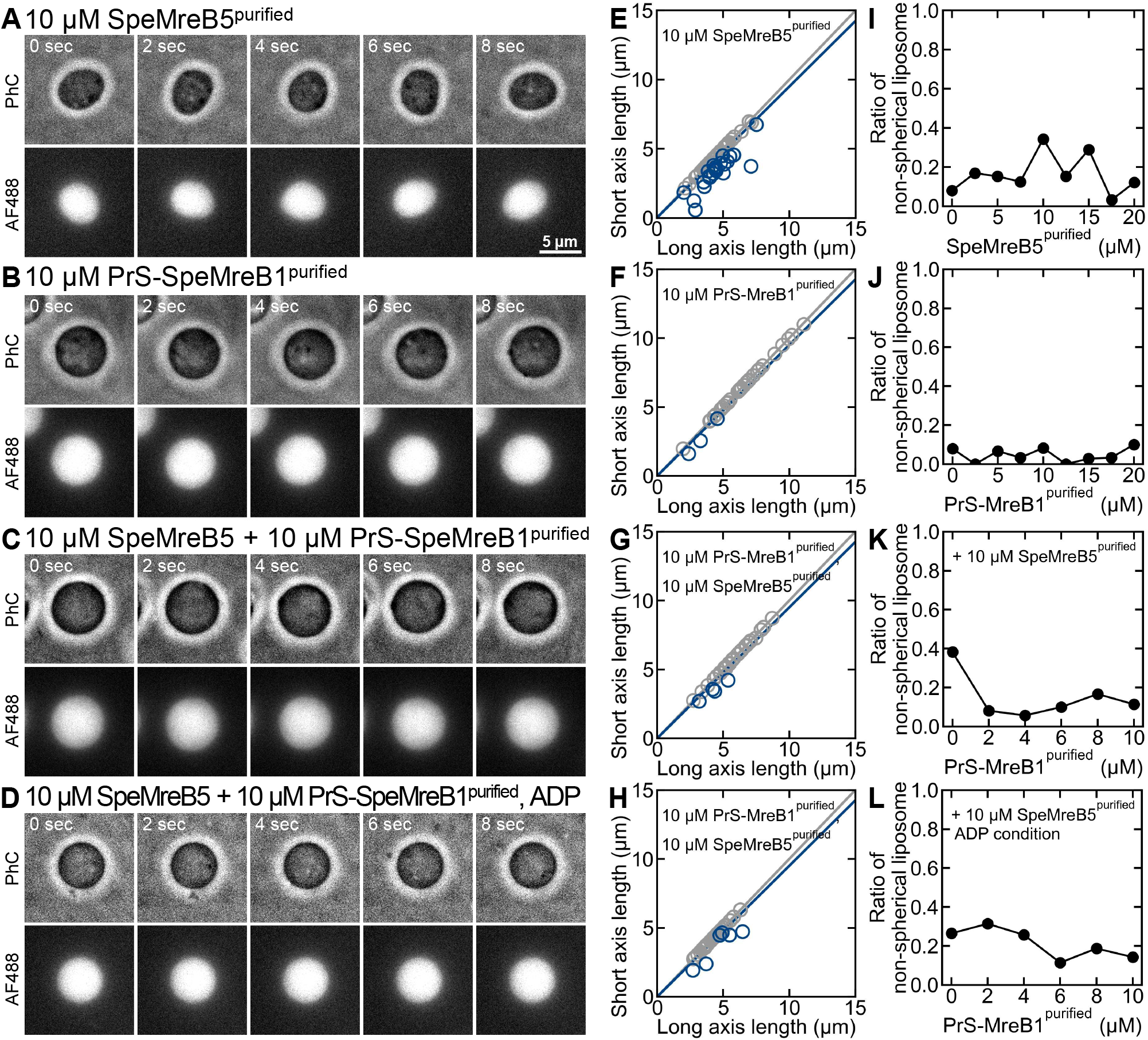
Purified SpeMreBs exert opposing effects on liposome deformation. (A–D) Representative time-lapse images of liposomes encapsulating SpeMreBs^purified^ **(see Movies S11–S15)**. The upper panels show phase-contrast images and the lower panels show Alexa Fluor 488 fluorescence images. Time stamps correspond to the phase-contrast images. Phase-contrast and fluorescence images were not acquired simultaneously. Fluorescence images were taken immediately after the phase-contrast time series. Scale bar, 5 μm. (E–H) Scatter plots of long-axis and short-axis lengths of individual liposomes under each condition. The black diagonal line indicates an aspect ratio of 1 (spherical liposomes). The blue line indicates an aspect ratio of 0.95. Liposomes with aspect ratios <0.95 were classified as deformed (blue), whereas those ≥ 0.95 were classified as spherical (black). (I–L) Fraction of deformed liposomes as a function of the type and concentration of encapsulated proteins. SpeMreB5^purified^ alone produced maximal deformation at ∼10 μM, with lower frequencies at higher concentrations (I). Liposomes encapsulating PrS-SpeMreB1^purified^ alone were typically larger but remained mostly spherical (J). When SpeMreB5^purified^ (10 μM) was combined with increasing concentrations of PrS-SpeMreB1, the fraction of deformed liposomes decreased (K). Replacement of ATP with ADP in the internal solution reduced the deformation-suppressing effect of PrS-SpeMreB1^purified^ (L).

SpeMreB5^purified^ alone induced concentration-dependent liposome deformation. Within the range of 0-10 μM, where filament formation has been observed by electron microscopy, the fraction of deformed liposomes increased with protein concentration **(Fig. 3E and Fig. S2A)**. Occasionally, elongated liposomes resembling the shape of *Spiroplasma eriocheiris* cells were observed **(Fig. S2B and Movie S12)**. However, the fraction of deformed liposomes decreased to ∼10% at higher concentrations (≥15 μM). This decrease may reflect changes in filament organization at higher SpeMreB5^purified^ concentrations, such as sheet or bundle formation, which could reduce interactions with the membrane. Both stable and fluctuating membrane behaviors were observed among the deformed liposomes under all SpeMreB5^purified^ conditions **(Fig. S2C)**.

In contrast, liposomes encapsulating PrS-SpeMreB1^purified^ alone remained largely spherical at all concentrations tested **(Fig. 3F and 3J)**. Although the size distribution shifted toward larger diameters, most liposomes remained spherical **(Fig. S2D)**. The few deformed liposomes that appeared were predominantly stable rather than fluctuating **(Fig. S2E)**. Notably, the frequency of deformation was even lower than that observed in liposomes containing control syn3B lysate lacking expressed SpeMreBs **(Fig. 1A)**, suggesting that PrS-SpeMreB1^purified^ interacts with the membrane in a way that stabilizes a larger, more spherical liposome shape.

We next examined whether PrS-SpeMreB1^purified^ influences SpeMreB5^purified^-induced deformation. When PrS-SpeMreB1^purified^ was mixed with 10 μM SpeMreB5^purified^, the condition that produced the highest deformation frequency, the fraction of deformed liposomes decreased markedly even with only 2 μM PrS-SpeMreB1^purified^. **(Fig. 3G and 3K)**. Analysis of aspect ratios confirmed that increasing concentrations of PrS–SpeMreB1^purified^ progressively suppressed liposome deformation **(Fig. S3A and S3C)**. Finally, we tested whether this antagonistic effect depends on the nucleotide state. When ATP in the internal solution was replaced with ADP, PrS-SpeMreB1^purified^ was less effective at restoring spherical liposome shapes **(Fig. 3H and 3L)**. The fraction of liposomes exhibiting membrane fluctuations increased **(Fig. S3C and S3D)**.

Together, these results indicate that SpeMreB5^purified^ actively promotes membrane deformation, whereas PrS-SpeMreB1^purified^ counteracts this activity and stabilizes spherical liposome shapes. The antagonistic interaction between the two proteins is influenced by the nucleotide state, consistent with the mechanism proposed by Takahashi et al. (2025) (20). These findings suggest that the *Spiroplasma* motility machinery contains opposing molecular activities that regulate filament stability and membrane deformation.

### Opposing effects of SpeMreB1 and SpeMreB5 on liposome morphology

To investigate the individual contributions of PrS-SpeMreB1^purified^ (SpeMreB4 class) and SpeMreB5^purified^, we titrated SpeMreB5^purified^ or PrS-SpeMreB1^purified^ into liposomes containing syn3B lysates expressing SpeMreB4 and SpeMreB5-mCherry **(Fig. 4)**. At low concentrations (2– 6 µM), SpeMreB5^purified^ did not alter the frequency of liposome deformation compared to lysate alone. However, at higher concentrations (10 µM), SpeMreB5^purified^ induced deformed liposome. A small subset of these liposomes exhibited spindle-shape, but such liposomes were rare and not analyzed quantitatively. Occasionally, bundles containing SpeMreB5^purified^ that aligned along the long axis were observed **(Fig. 4A and Movie S16)**. Despite these structural changes, SpeMreB5^purified^ did not show the extreme aspect ratios characteristic of *Spiroplasma* **(Fig. 4B)**. Furthermore, the overall frequency of deformed liposomes remained constant **(Figs.s 4C and S4A)**. These observations suggest that SpeMreB5^syn3B lysate^ may already be near saturation, or that additional factors required for further deformation are limited. In contrast, the addition of PrS-SpeMreB1^purified^ promoted a spherical shape **(Fig. 4D and Movie S17)**. Higher concentrations of PrS-SpeMreB1^purified^ led to a dose-dependent increase in spherical liposomes and a reciprocal decrease in deformed populations **(Figs. 4E, 4F, and S4B)**. These results reinforce our model that SpeMreB1/4 antagonizes SpeMreB5-induced membrane deformation, driving the membrane toward a spherical shape. We noted that fluctuating membranes were frequently observed. However, the proportion of stable, non-spherical shapes did not increase significantly **(Figs. S4C and S4D)**. This dynamic behavior may be due to the increased salt concentration (+150 mM) required for purified proteins.

**Figure 4.**
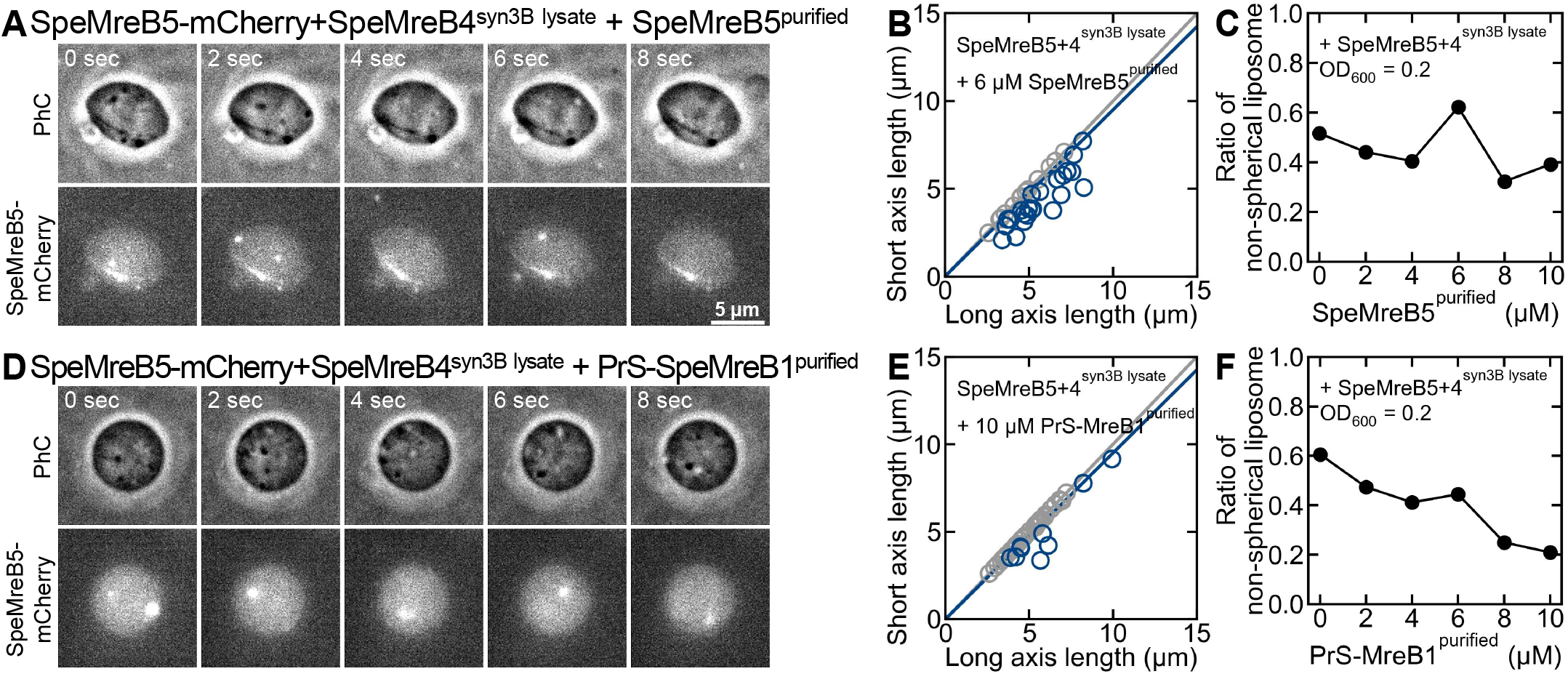
SpeMreB1 and SpeMreB5 reciprocally regulate liposome morphology. (A) Representative phase-contrast (upper) and fluorescence (lower) images of liposomes containing SpeMreB5-mCherry/SpeMreB4^syn3B lysate^ in the presence of 10 µM SpeMreB5^purified^ **(see Movie S16)**. High SpeMreB5^purified^ concentrations induced spindle-shape liposome with internal bundles aligned along the long axis. Scale bar, 5 µm. (B) Distribution of long and short axes for liposomes under conditions in (A). Extreme aspect ratios were not detected, and no clear concentration dependence was observed. (C) Frequency of deformed liposomes as a function of SpeMreB5^purified^ concentration. (D) Representative images of liposomes containing SpeMreB5-mCherry/SpeMreB4^syn3B lysate^ in the presence of 10 µM PrS-SpeMreB1^purified^ **(see Movie S17)**. PrS-SpeMreB1^purified^ addition shifted the population toward spherical. (E) Distribution of long and short axes for liposomes in the presence of PrS-SpeMreB1^purified^. (F) Frequency of non-spherical liposomes across varying PrS-SpeMreB1^purified^ concentrations. SpeMreB1^purified^ dose-dependently inhibited SpeMreB5^purified^ -induced membrane deformation.

### SpeMreB4^syn3B lysate^ and ATP depletion modulate SpeMreB5^syn3B lysate^ particle dynamics in liposomes

To characterize the behavior of SpeMreBs within the liposome, we performed particle-tracking analysis using syn3B lysates expressing SpeMreB5-mCherry^syn3B lysate^ as a fluorescent marker **(Fig. 5A and Movie S18)**. We monitored the displacements of SpeMreB5-mCherry^syn3B lysate^ particles for up to 10 seconds within spherical liposomes (4-5 µm diameter) at a constant lysate concentration **(Figs. 5B–5D)**. Particles moved randomly within the central region of the liposome or along the membrane **(Fig. S5)**. We terminated tracking when particles moved out of the focal plane. The mobility of SpeMreB5-mCherry^syn3B lysate^ depended on the presence of SpeMreB4^syn3B lysate^ and the availability of ATP. In liposomes containing only SpeMreB5-mCherry^syn3B lysate^, the diffusion coefficient (*D*) was 1.3 ± 0.74 µm^2^/s **(Fig. 5E)**. Co-expression of SpeMreB4^syn3B lysate^ slightly increased this mobility to D = 2.0 ± 2.2 µm^2^/s **(Fig. 5F)**. Notably, ATP depletion via hexokinase addition further enhanced particle displacement, increasing the diffusion coefficient to 3.4 ± 4.4 µm^2^/s **(Fig. 5G)**. These results show that SpeMreB4^syn3B lysate^ and ATP levels collectively regulate SpeMreB5^syn3B lysate^ particle dynamics. The increased particle mobility upon ATP depletion suggests that SpeMreBs interactions within these clusters are influenced by ATP.

**Figure 5.**
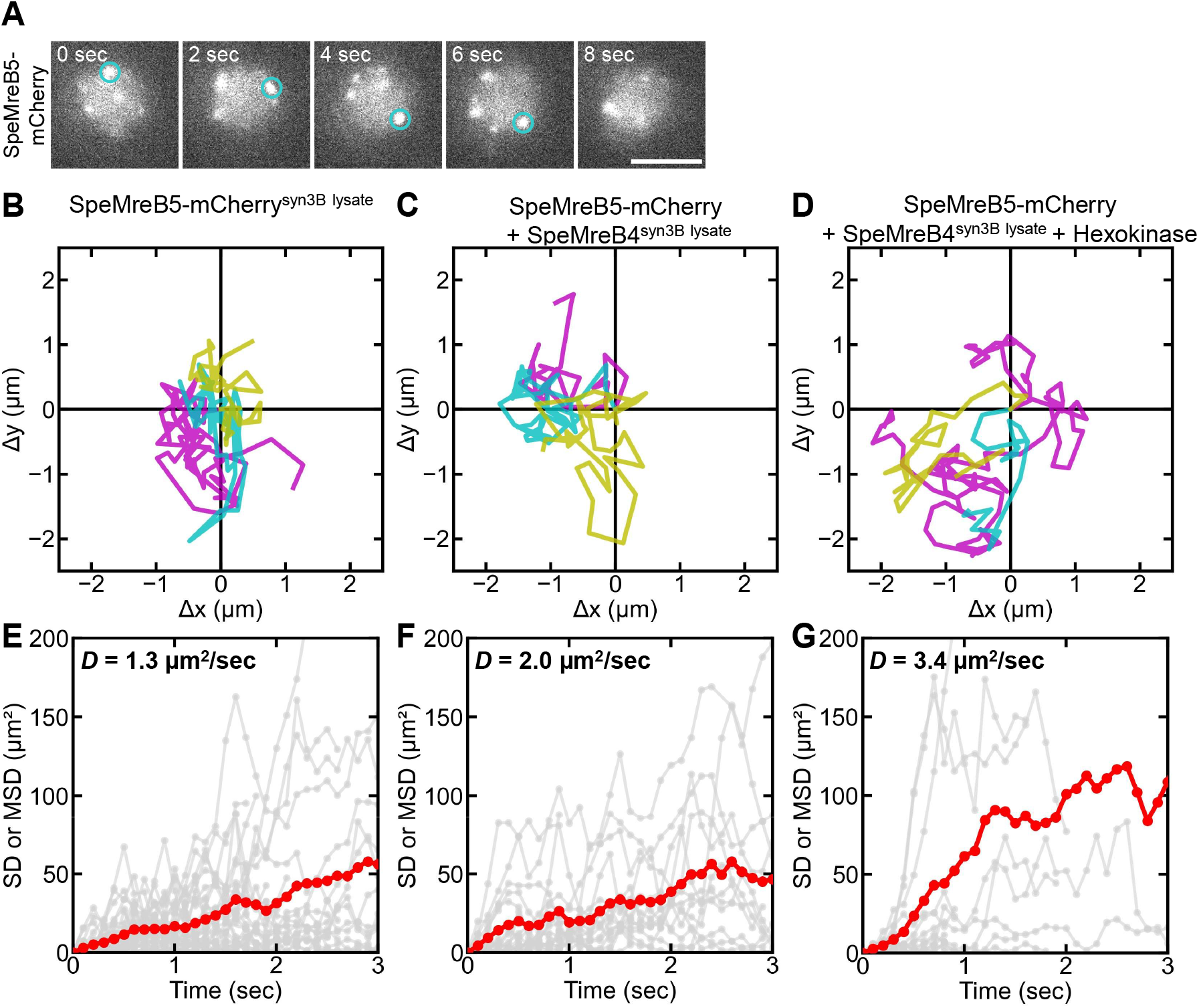
SpeMreB4 and ATP depletion modulate the dynamics of SpeMreB5^syn3B lysate^ particles. (A) Fluorescence images of SpeMreB5-mCherry^syn3B lysate^ particles within a liposome **(see Movie S18)**. Circles indicate a tracked particle corresponding to one of the traces in (B, blue). Scale bar, 5 µm. (B–D) Representative trajectories of SpeMreB5-mCherry^syn3B lysate^ particles in liposomes of similar size (OD_600_ = 0.2). Displacements were tracked at 0.1 s intervals for up to 10 s. Tracking ended when particles left the focal plane. Each condition was shown above each panel. (E–G) Mean squared displacement (MSD) analysis. Individual MSD curves (gray) and the mean MSD (red) are shown. Diffusion coefficients (*D*) were estimated from the first 1 s of observation. (E) SpeMreB5-mCherry^syn3B lysate^ alone (*D* = 1.3 µm^2^/s, n=34). (F) Lysate co-expressing SpeMreB5-mCherry^syn3B lysate^ and SpeMreB4^syn3B lysate^ (*D* =2.0 µm^2^/s, n = 18). (G) ATP-depleted lysate containing SpeMreB5-mCherry^syn3B lysate^ and SpeMreB4^syn3B lysate^ supplemented with hexokinase (*D* = 3.4 µm^2^/s, n = 13).

## Discussion

### Principles of SpeMreB-driven liposome deformation

Encapsulating SpeMreBs from syn3B lysates or purified *E. coli* systems into liposomes did not reproduce the helical geometry or swimming motility of *Spiroplasma*. Thus, while SpeMreBs are essential components, helical deformation likely requires a specific cellular context established during cell growth, such as defined membrane composition or local protein density, which may not be recapitulated in the reconstituted system. Nevertheless, our reconstitution assay found three fundamental principles of membrane deformation (i) SpeMreB5 autonomously induces liposome deformation **(Fig. 6A, left)**, (ii) SpeMreB1 (and SpeMreB4) suppresses this deformation **(Fig. 6A, right)**, and (iii) these effects are governed by the nucleotide state.

**Figure 6.**
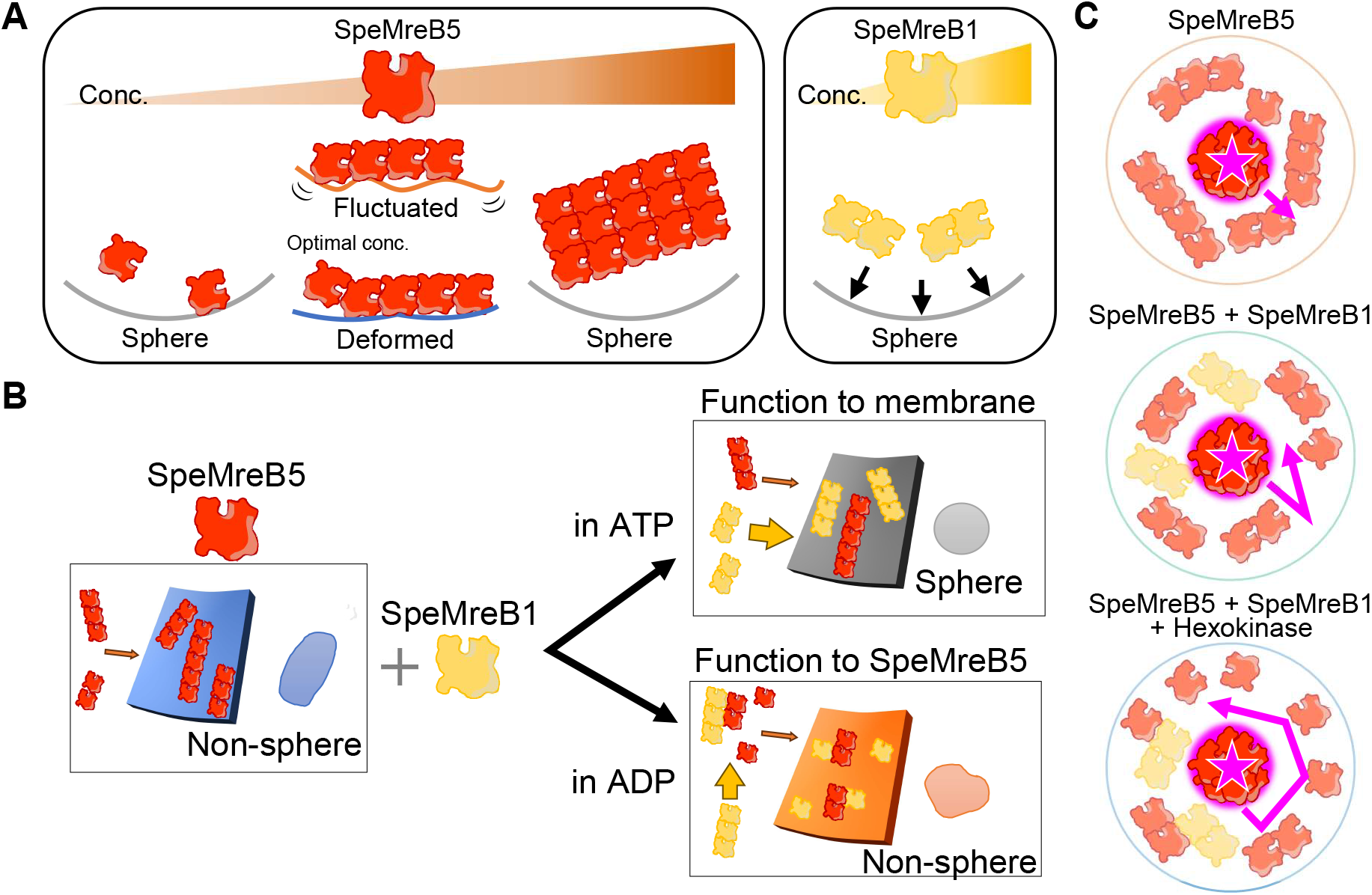
Model of reciprocal membrane regulation by SpeMreB5 and SpeMreB1/4. (A) (Left) SpeMreB5 changes mechanical stresses on membranes. At optimal concentrations, it promotes stable non-spherical geometries, whereas excessive concentrations cause localized bundling that impairs uniform membrane support. (Right) SpeMreB1/4 promotes a spherical shape across all tested concentrations. (B) Mechanistic interplay between SpeMreBs and nucleotides. Arrows indicate the direction and relative strength of interactions. In the presence of ATP, SpeMreB5 drives deformation, while SpeMreB1/4 antagonizes this process presumably through direct membrane interaction. In the presence of ADP, the spherical-restoring effect of SpeMreB1/4 is weakened because of reduced membrane affinity, despite its enhanced ability to destabilize SpeMreB5 filaments. (C) Internal cytoskeletal stability inferred from particle tracking. (Top) Without SpeMreB4, SpeMreB5 forms a high-order network that restricts particle diffusion. (Middle/Bottom). Addition of SpeMreB4 or ATP depletion destabilizes this network; consequently, the diffusion coefficient (*D*) increases as the internal architecture becomes more destabilized.

### Mechanical antagonism and nucleotide regulation

In liposomes containing only SpeMreB5, the proportion of deformed liposomes increased in a concentration-dependent manner. Although the magnitude of the effect differed between syn3B lysates and purified proteins, these data show that SpeMreB5 can exert force on the membrane. While the membrane-binding affinity of SpeMreB5 is relatively weak and its nucleotide dependence is limited **(Figs. 3I and 3L)**, the addition of SpeMreB1/4 significantly suppressed the appearance of deformed liposomes **(Figs. 1G and 3K)**. This inhibitory effect is consistent with the previously reported ability of SpeMreB1/4 to destabilize high-order SpeMreB5 structures (20).

### Nucleotide-dependent regulation of liposome shape

Under ADP conditions, the suppression of deformation by PrS-SpeMreB1^purified^ was weakened **(Fig. 3L)**. Similarly, deformed liposomes persisted in syn3B lysate upon ATP depletion via hexokinase **(Fig. 2B)**. If deformation were determined simply by the amount of SpeMreB filaments, the enhanced destabilizing effect of SpeMreB1/4 in the ADP state should have further reduced the number of deformed liposomes. Our results suggest that liposome deformation is not solely determined by the quantity of SpeMreB assemblies but is also influenced by the interaction between SpeMreBs and the membrane. We propose a model where SpeMreB1/4 activity involves at least two parameters, (i) destabilization of high-order SpeMreB5 structures and (ii) membrane-binding affinity. Assuming that SpeMreB1/4 interacts more strongly with the membrane in the presence of ATP, it may antagonize SpeMreB5-induced deformation through membrane binding, thereby restoring a spherical shape. In the ADP-bound state, the destabilizing effect on high-order SpeMreB5 structures may increase, but reduced membrane affinity would explain the weakened spherical-restoring effect **(Fig. 6B)**. This model remains a hypothesis based on functional correlations, as protein membrane affinity and nucleotide occupancy were not quantified in this study.

### Internal dynamics as a readout of cytoskeletal stability

Particle-tracking analysis showed that SpeMreB5-mCherry^syn3B lysate^ clusters diffuse in the central region of the liposome. These particles did not localize specifically to deformation sites and were primarily observed in the central region, consistent with reports that filamentous structures tend to occupy the periphery while particulate matter remains central (28). The increased diffusion coefficients (*D*) observed upon SpeMreB4^syn3B lysate^ expression or ATP depletion **(Figs. 5D and 5G)** may reflect the destabilization of the internal high-order network. The particularly high mobility observed under ATP depletion confirms that nucleotide hydrolysis is an important regulator of internal scaffold stability. The direct link between particle mobility and liposome deformation is not yet clear. However, these dynamics provide a readout of the cytoskeletal state as regulated by ATP hydrolysis.

In conclusion, SpeMreB5 serves as a factor capable of changing the mechanical stresses on membranes, while SpeMreB1/4 modulates it by altering both filament stability and membrane interaction in a nucleotide-dependent manner. SpeMreB1/4 does not act as a simple inhibitor but rather as an actuator whose activity shifts with the ATP state. It seems that the antagonistic interactions between these two SpeMreBs on the membrane surface are actively causing the membrane fluctuations observed. The failure to reproduce helical motility suggests that spatial constraints and membrane composition are critical. High-speed swimming likely requires coordinated, transient interactions between SpeMreB4 and SpeMreB5, which are favored by the close proximity of proteins within the native cellular environment.

## Materials & Methods

### Materials

1,2-dioleoyl-*sn*-glycero-3-phosphatidylcholine (DOPC), was purchased from Fujifilm Wako Pure Chemical Industries (Osaka, Japan). AF 488 NHS ester was purchased from Lumiprobe (Hallandale Beach, FL, USA). ATP, EGTA, and all salts were of analytical grade and were purchased from Fujifilm Wako Pure Chemical Industries. All chemicals were used without further purification. Free Alexa Fluor 488 dye in the internal solution was prepared from excess dye remaining after protein-labeling reactions.

### Bacterial strains

JCVI-syn3B (GenBank, CP069345.1), and syn3B strains expressing SpeMreB5-mCherry-, or SpeMreB5-mCherry-SpeMreB4-expressing were used as described previously (1). Expression of SpeMreB1 and SpeMreB5 in syn3B resulted in reduced motility compared to SpeMreB4 and SpeMreB5 and was not used further. Cells were cultured in SP4 medium (18, 29) at 37°C. After 2 days of growth, cells were harvested by centrifugation at 10,000 × g for 10 minutes at 4°C, washed twice with PBS, and collected by centrifugation under the same conditions. Cell pellets were stored at -80°C until use.

### Protein purification

PrS-SpeMreB1 and SpeMreB5 were purified by Ni–NTA affinity chromatography followed by gel filtration as described previously (19, 30, 31). Briefly, cell pellets were resuspended in His trap buffer A (50 mM Tris–HCl (pH 8.0), 300 mM NaCl, 50 mM imidazole–HCl (pH 8.0)), centrifuged at 12,000 x g for 30 min at 4 °C. The supernatant was applied to 5 ml Ni-NTA resin (Fujifilm). PrS-SpeMreB1 was eluted with 500 mM imidazole because it contains tandem 6× His tags, whereas SpeMreB5 was eluted with 230 mM imidazole. Purified SpeMreBs were stored in 50 mM Tris-HCl (pH8.0) and 200 mM NaCl on ice. When aggregations were observed during the storage, samples were centrifuged at 55,000 × g for 20 min at 4°C, and the supernatant was used. Protein concentrations were determined from the absorbance at 280 nm measured using a NanoDrop One (Thermo Fisher Scientific), with the absorption coefficients estimated from ProtParam (31). The protein (PrS) tag on SepMreB1 was retained to improved solubility.

### Liposome

Liposomes were prepared using a reduced volume W/O emulsion centrifugation method (32). All procedures were performed at room temperature unless otherwise stated. Frozen syn3B cell pellets were lysed by ultrasonication in Internal Solution buffer (IS, 20 mM imidazole (pH 7.0, 100 mM KCl, 2 mM MgCl_2_, 2 mM ATP, 0.5 mM EGTA, 500 mM sucrose, 1 mg/mL BSA, 16 µM AF 488). Lysate concentrations were adjusted with Internal buffer based on OD_600_ measurement immediately after lysis. DOPC was dissolved in liquid paraffin at a final concentration of 1 mM by incubation at 60°C for 30 min. Lipid solution of 60 µL was mixed with IS buffer of 1.5 or 3 µL containing the sample by vigorous tapping to form a W/O emulsion in 0.6 ml test tube. The emulsion was layered onto 60 µL of Outer Solution buffer (OS, 20 mM Imidazole (pH 7.0), 100 mM KCl, 2 mM MgCl_2_, 2 mM ATP, 0.5 mM EGTA, 500 mM glucose, 1 mg/mL BSA) in a PCR tube and centrifuged at 10,000 × g for 10 min at 4°C. After centrifugation, liposomes accumulated at the bottom of the tube. The oil phase and approximately 50 µL of OS buffer were removed, and the pellet was gently suspended in 60–120 µL of fresh OS buffer.

For ATP depletion experiments, 0.05 units/µL of hexokinase and 100 mM glucose were added to the IS buffer, and an additional 100 mM glucose was included in the OS buffer. When purified SpeMreBs were encapsulated, OS buffer was supplemented with Tris-HCl (pH 8.0) and NaCl to a final concentration of 30.5 mM and 122 mM, respectively, to compensate for ions carried with the protein stock.

### Image Acquisition and Analysis

Liposomes were observed at room temperature using phase contrast and fluorescence microscopes (ECLIPSE Ti2-E (Nikon, Tokyo, Japan)) equipped with a 100× objective lens and 488 and 561 nm lasers. Images were acquired with ORCA-Fusion camera (HAMAMATSU, Japan) using 70 msec exposure times, corresponding to a pixel size of 0.06 µm in the sample plane. Time lapse images were recorded at 10 frames/s for 10 sec over a field of view of 1192 × 1192 pixels (∼77 × 77 µm). For long term observation, one frame was acquired every 5 min up to 12 h.

Coverslips were cleaned by sonication in KOH and ultrapure water, air-dried and assembled into flow chambers using double sided tape and a slide glass. Liposomes were loaded into the chamber, which was sealed with white petrolatum to prevent evaporation. After sealing, chambers were equilibrated for 10 min before imaging Approximately 10 liposomes were typically observed per chamber, and more than 30 liposomes were analyzed for each experimental condition.

Liposome axial length was measured using ImageJ (NIH). Only isolated single liposomes were analyzed; aggregates were excluded. Encapsulation of IS buffer was verified by comparing fluorescence and phase contrast images. For non-spherical liposomes, the axial length was measured at the point showing the maximum aspect ratio. Particle tracking analysis was performed using the ImageJ plugin SpeckleTrackerJ, developed by Dr. Dimitrios Vavylosnis, Lehigh University), following instructions from Dr. Sawako Yamashiro, Kyoto University (33). For quantitative comparison of Brownian motion, analyses were restricted to liposomes with diameters of 4–5 μm, and lysate concentrations were standardized to OD_600_ = 0.2.

## Acknowledgments

We thank Dr. Daichi Takahashi (Okayama University) for providing plasmids and for discussions on SpeMreB function for *Spiroplasma* swimming. We appreciate Dr. Hideaki Matsubayashi (Tohoku University) for helpful discussions regarding lipid components. We also thank Ms. Mone Mimura and Mr. Satoshi Kanamori (Osaka Metropolitan University) for discussions regarding the swimming mechanism. We are grateful to Drs. Shigeyuki Kakizawa (AIST) Masaki Mizutani (Gakushuin University) for providing syn3B culture protocols, and to Dr. Yoshihiro Yamaguchi (Osaka Metropolitan University) for technical assistance with SpeMreB construction and the pCold-PrS expression system. We thank Prof. Robert C. Robinson (Okayama University and VISTEC) for suggesting the ATP depletion approach.

This study was supported by JKA (24MC1001-094), the Union Tool Foundation, the Japan Society for the Promotion of Science (JSPS) KAKENHI Grant Number (24K08218), and JSPS Program for Forming Japan’s Peak Research Universities(J-PEAKS, JPJS00420240017) to I.F.; NUT-SPRING (JPMJSP2189) to T.M.; and JST CREST Grant Number JPMJCR19S5 to M.M. H.K. is a recipient of a JSPS Research Fellowship (JP24KJ0189).

## Author Contributions

T.M., M.M., and I.F. conceived the study and designed the experiments. T.M., M.H., and K.T. developed the methodology. H.K. and A.A. prepared the syn3B strains. T.M. and T.N. performed the investigations and visualized the data. M.H., K.T., M.M., and I.F. supervised the research. T.M., M.H., K.T., M.M., and I.F. wrote and edited the manuscript. All authors have given approval to the final version of the manuscript.

