## Supplementary material for "Coordinated Membrane Deformation Driven by a Minimal Set of *Spiroplasma* MreB Isoforms": SI figures and legends

### Sup. Figure legends for

**Fig. S1. Deformation of liposomes containing SpeMreBs$^{\text{syn3B lysates}}$.**
(A–C) Long and short axis lengths of liposomes containing (A) original syn3B lysate, (B) syn3B lysate expressing SpeMreB5-mCherry, or (C) syn3B lysate co-expressing SpeMreB5-mCherry and SpeMreB4. Lysate concentrations ($OD_{600}$) are indicated in each plot. Gray lines represent an aspect ratio (short/long axis) of 1.0 (spheres); blue lines represent an aspect ratio of 0.95. Blue circles denote deformed liposomes (aspect ratio < 0.95); gray circles denote spherical liposomes (aspect ratio ≥ 0.95). (D) Aspect ratios of all observed liposomes (n = 1452). Bin size is 0.065 (μm/μm).

**Fig. S2. Deformation of liposomes containing SpeMreB$^{\text{purified}}$.**
(A) Aspect ratios and dimensions of liposomes containing SpeMreB5$^{\text{purified}}$. Symbols and lines are as in Fig. S1. (B) Elongated liposomes containing 8 μM SpeMreB5$^{\text{purified}}$. Upper: Phase-contrast; Lower: AF488 fluorescence. Time is shown in phase-contrast images. Scale bar (bottom right), 5 µm. (C) Concentration-dependent liposome behavior of SpeMreB5$^{\text{purified}}$-containing liposomes. Gray: Sphere; Orange: Fluctuated; Blue: Deformed. (D) Aspect ratios of liposomes containing purified PrS-SpeMreB1$^{\text{purified}}$. Symbols are as in (A). (E) Fraction of behavior of PrS-SpeMreB1$^{\text{purified}}$-containing liposomes. Colors are as in (C).

**Fig. S3. Deformation of liposomes containing SpeMreB5$^{\text{purified}}$ and PrS-SpeMreB1$^{\text{purified}}$.**
(A, B) Aspect ratios of liposomes containing SpeMreB5$^{\text{purified}}$ and PrS-SpeMreB1$^{\text{purified}}$ in the presence of (A) ATP or (B) ADP. Symbols and lines are as in Fig. S1. (C, D) Fraction of liposome behaviors for (C) ATP or (D) ADP conditions. Colors are as in Fig. S2.

**Fig. S4. Effect of additional PrS-SpeMreB1$^{\text{purified}}$ or SpeMreB5$^{\text{purified}}$ on SpeMreBs$^{\text{syn3B lysate}}$ containing liposomes.**
(A, B) Aspect ratios of liposomes containing SpeMreB5-mCherry/SpeMreB4$^{\text{syn3B lysate}}$ additionally in the presence of the indicated concentration of (A) SpeMreB5$^{\text{purified}}$ or (B) PrS-SpeMreB1$^{\text{purified}}$. Symbols and lines are as in Fig. S1. (C, D) Fraction of liposome behaviors additionally in the presence of (C) SpeMreB5$^{\text{purified}}$ or (D) PrS-SpeMreB1$^{\text{purified}}$. Colors are as in Fig. S2.

**Fig. S5. Trajectories of SpeMreB5–mCherry$^{\text{syn3B lysate}}$ particles within liposomes.**
(A–C) Representative trajectories of SpeMreB5-mCherry$^{\text{syn3B lysate}}$ particles in syn3B lysates expressing (A) SpeMreB5-mCherry alone, (B) SpeMreB5-mCherry and SpeMreB4, or (C) SpeMreB5-mCherry and SpeMreB4 with hexokinase. Trajectories in (A) and (B) (left columns) exhibit transient movement along the membrane. Observation times: (A) 3.6, 3.5, 4.1 s; (B) 4.7, 2.9, 3.9 s; (C) 1.6, 9.7, 2.0 s. Particle positions were recorded at 0.1 s intervals.

Sup. Movie legend

**Mov. S1. Phase-contrast images of liposomes containing original syn3B lysate.**
Original syn3B lysate ($OD_{600}$ = 0.3). White arrows: spherical liposomes; yellow arrows: non-spherical liposomes. Most liposomes remain spherical without SpeMreB expression. 10 s, 10 fps. Scale bar, 5 μm.

**Mov. S2. Representative liposome containing original syn3B lysate.**
Left: Phase-contrast; Right: AF488 fluorescence. 10 s, 10 fps. Scale bar, 5 μm.

**Mov. S3. Phase-contrast images of liposomes containing SpeMreB5-expressing syn3B lysate.**
syn3B MreB5-mCherry lysate ($OD_{600}$ = 0.3). White arrows: spherical; blue arrows: stable deformed; yellow arrows: fluctuating membranes. Deformation increases with lysate concentration. 10 s, 10 fps. Scale bar, 5 μm.

**Mov. S4. Deformation of liposome containing SpeMreB5-expressing syn3B lysate.**
Left: Phase-contrast; Right: SpeMreB5-mCherry fluorescence. 10 s, 10 fps. Scale bar, 5 μm.

**Mov. S5. Phase-contrast images of liposomes containing SpeMreB5/SpeMreB4 co-expressing syn3B lysate.**
Liposomes containing SpeMreB5-mCherry$^{\text{syn3B lysate}}$/SpeMreB4$^{\text{syn3B lysate}}$ ($OD_{600}$ = 0.3). White arrows: spherical; other colors: non-spherical. SpeMreB4 reduces the frequency of deformation compared to SpeMreB5 alone. 10 s, 10 fps. Scale bar, 5 μm.

**Mov. S6. Deformation of liposomes containing SpeMreB5/MreB4 co-expressing syn3B lysate.**
Left: Phase-contrast; Right: SpeMreB5-mCherry fluorescence. 10 s, 10 fps. Scale bar, 5 μm.

**Mov. S7. Non-spherical liposome maintaining a stable shape.**
Liposome containing SpeMreB5-mCherry$^{\text{syn3B lysate}}$/SpeMreB4$^{\text{syn3B lysate}}$ ($OD_{600}$ = 0.1). The deformed shape is maintained during rotation. Left: Phase-contrast; Center: SpeMreB5-mCherry; Right: AF488. 10 s, 10 fps. Scale bar, 5 μm.

**Mov.S8. Fluctuating liposome.**
Liposome containing SpeMreB5-mCherry$^{\text{syn3B lysate}}$/SpeMreB4$^{\text{syn3B lysate}}$ ($OD_{600}$ = 0.1). The membrane fluctuates while moving. Left: Phase-contrast; Center: SpeMreB5-mCherry; Right: AF488. 10 s, 10 fps. Scale bar, 5 μm.

**Mov. S9. Long-term observation of deformed liposome.**
Liposome containing MreB5-mCherry$^{\text{syn3B lysate}}$/SpeMreB4$^{\text{syn3B lysate}}$ ($OD_{600}$ = 0.5) with hexokinase. Recorded for 12 h (5 min/frame). Time is shown at top left. 10 fps. Scale bar, 5 μm.

**Mov. S10. Deformed liposomes returning to a spherical shape.**
Conditions as in Mov. S9. White arrow: recovery of spherical shape; yellow arrow: rupture. Recorded for 12 h (5 min/frame). 10 fps speed. Scale bar, 5 μm.

**Mov. S11. Deformation of liposome containing SpeMreB5$^{\text{purified}}$.**
10 μM SpeMreB5$^{\text{purified}}$. Left: Phase-contrast; Right: AF488. 10 s, 10 fps. Scale bar, 5 μm.

**Mov. S12. String-like liposome containing 8 μM SpeMreB5$^{\text{purified}}$.**
Long, thin morphology resembling *Spiroplasma*. Left: Phase-contrast; Right: AF488. 10 s, 10 fps. Scale bar, 5 μm.

**Mov. S13. Liposome containing PrS-SpeMreB1$^{\text{purified}}$.**
Most liposomes containing 10 μM PrS-SpeMreB1$^{\text{purified}}$ remain spherical as this representative liposome. Left: Phase-contrast; Right: AF488. 10 s, 10 fps. Scale bar, 5 μm.

**Mov. S14. Reduced deformation in liposome containing SpeMreB5$^{\text{purified}}$ and PrS-SpeMreB1$^{\text{purified}}$.**

Liposome containing 10 µM SpeMreB5$^{\text{purified}}$ and 10 µM PrS-SpeMreB1$^{\text{purified}}$. Left: Phase-contrast; Right: AF488. 10 s, 10 fps. Scale bar, 5 µm.

**Mov. S15. Weakened effect of PrS-SpeMreB1$^{\text{purified}}$ under ADP conditions.**
Liposome containing 10 µM SpeMreB5$^{\text{purified}}$ and 10 µM PrS-SpeMreB1$^{\text{purified}}$ in ADP buffer. The frequency of non-spherical liposomes decreased more gradually in a PrS-SpeMreB1$^{\text{purified}}$ concentration-dependent manner. Left: Phase-contrast; Right: AF488. 10 s, 10 fps. Scale bar, 5 µm.

**Mov. S16. Effect of adding SpeMreB5$^{\text{purified}}$ to syn3B lysate.**
Liposome containing SpeMreB5-mCherry/SpeMreB4$^{\text{syn3B lysate}}$ ($OD_{600}$ = 0.2) in addition to 10 µM SpeMreB5$^{\text{purified}}$. SpeMreB5 bundles are visible in mCherry fluorescence. 10 s, 10 fps. Scale bar, 5 µm.

**Mov. S17. Effect of adding purified SpeMreB1 to syn3B lysate.**
SpeMreB5-mCherry/SpeMreB4$^{\text{syn3B lysate}}$ ($OD_{600}$ = 0.2) with 10 µM PrS-SpeMreB1$^{\text{purified}}$. 10 s, 10 fps. Scale bar, 5 µm. Non-spherical liposomes were decreased as increasing PrS-SpeMreB1 $^{\text{purified}}$.

**Mov. S18. Tracking of SpeMreB5-mCherry particles.**
Three moving particles of SpeMreB5-mCherry, observed within a syn3B lysate co-expressing SpeMreB5 and SpeMreB4, are highlighted with cyan circles. 10 s, 10 fps. Scale bar, 5 µm.

### Fig. S1

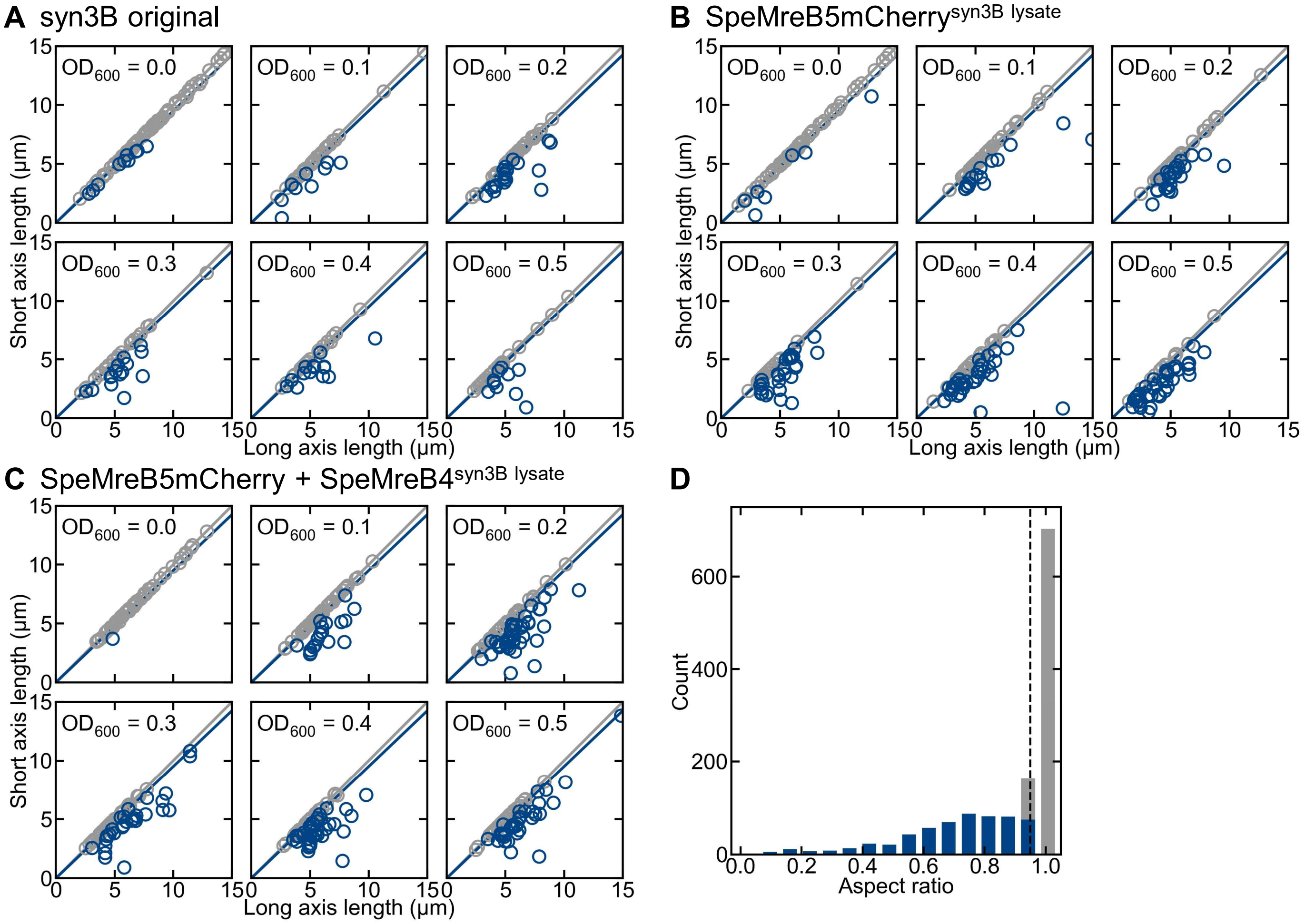
A syn3B original
B SpeMreB5mCherry^syn3B lysate
C SpeMreB5mCherry + SpeMreB4^syn3B lysate
D
OD600 = 0.0
OD600 = 0.1
OD600 = 0.2
OD600 = 0.3
OD600 = 0.4
OD600 = 0.5
Short axis length (μm)
Long axis length (μm)
Count
Aspect ratio

Fig. S2

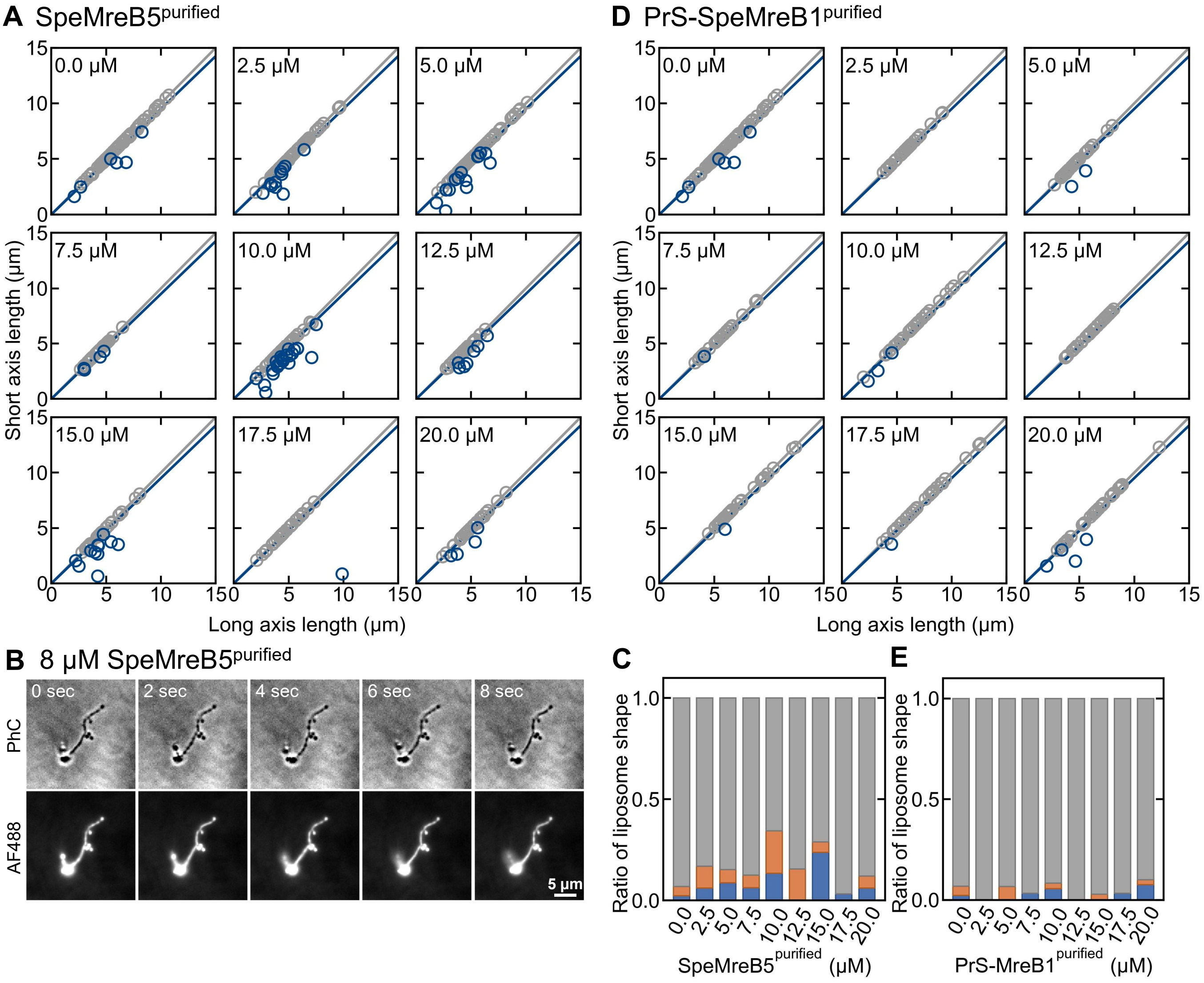

A SpeMreB5purified
D PrS-SpeMreB1purified
0.0 μM
2.5 μM
5.0 μM
7.5 μM
10.0 μM
12.5 μM
15.0 μM
17.5 μM
20.0 μM
Short axis length (μm)
Long axis length (μm)
0
5
10
15
B 8 μM SpeMreB5purified
0 sec
2 sec
4 sec
6 sec
8 sec
PhC
AF488
5 μm
C
E
Ratio of liposome shape
0.0
0.5
1.0
0.0
2.5
5.0
7.5
10.0
12.5
15.0
17.5
20.0
SpeMreB5purified (μM)
PrS-MreB1purified (μM)

Fig. S3

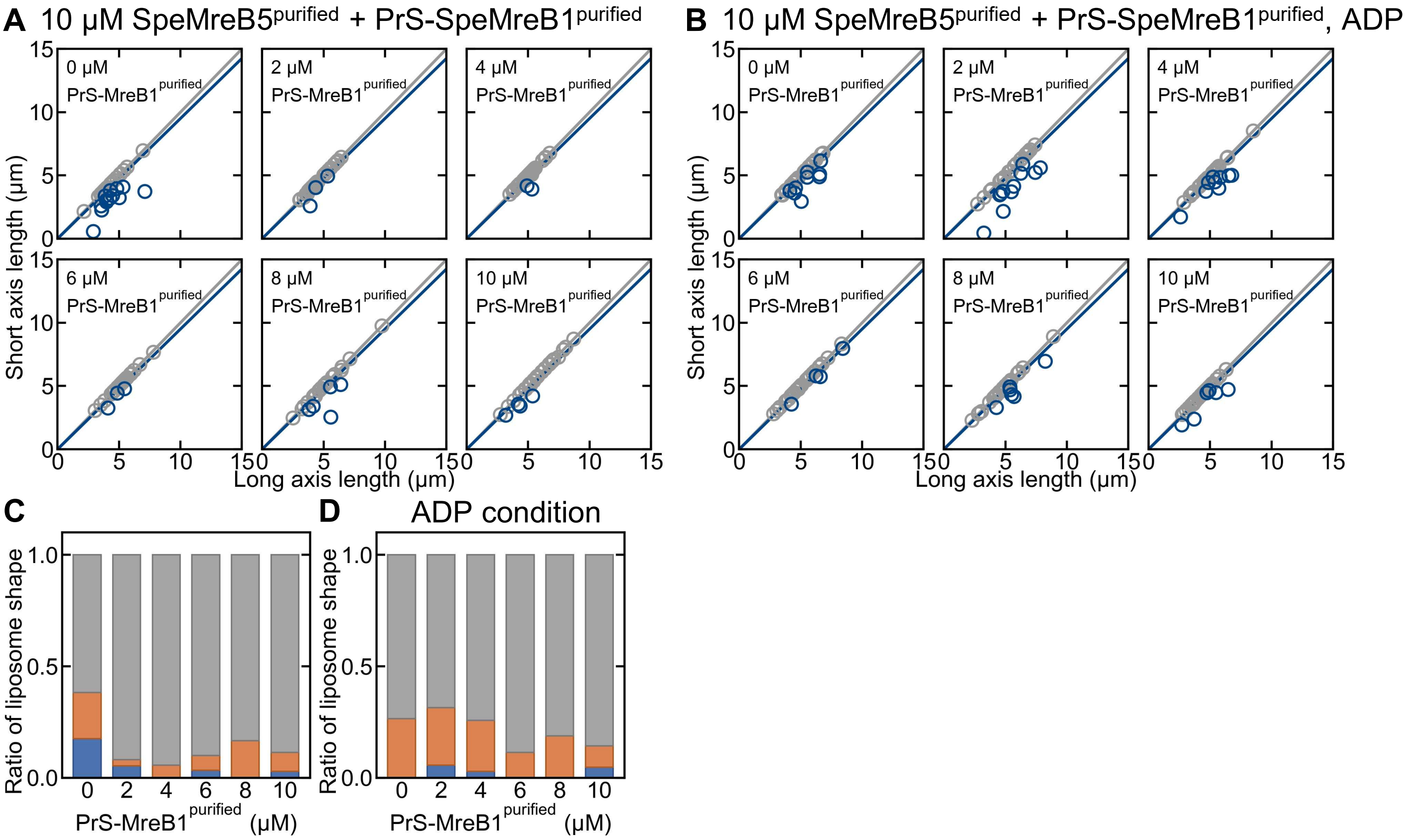
A 10 μM SpeMreB5^purified + PrS-SpeMreB1^purified
B 10 μM SpeMreB5^purified + PrS-SpeMreB1^purified, ADP
0 μM PrS-MreB1^purified
2 μM PrS-MreB1^purified
4 μM PrS-MreB1^purified
6 μM PrS-MreB1^purified
8 μM PrS-MreB1^purified
10 μM PrS-MreB1^purified
Short axis length (μm)
Long axis length (μm)
C
D ADP condition
Ratio of liposome shape
PrS-MreB1^purified (μM)

### Fig. S4

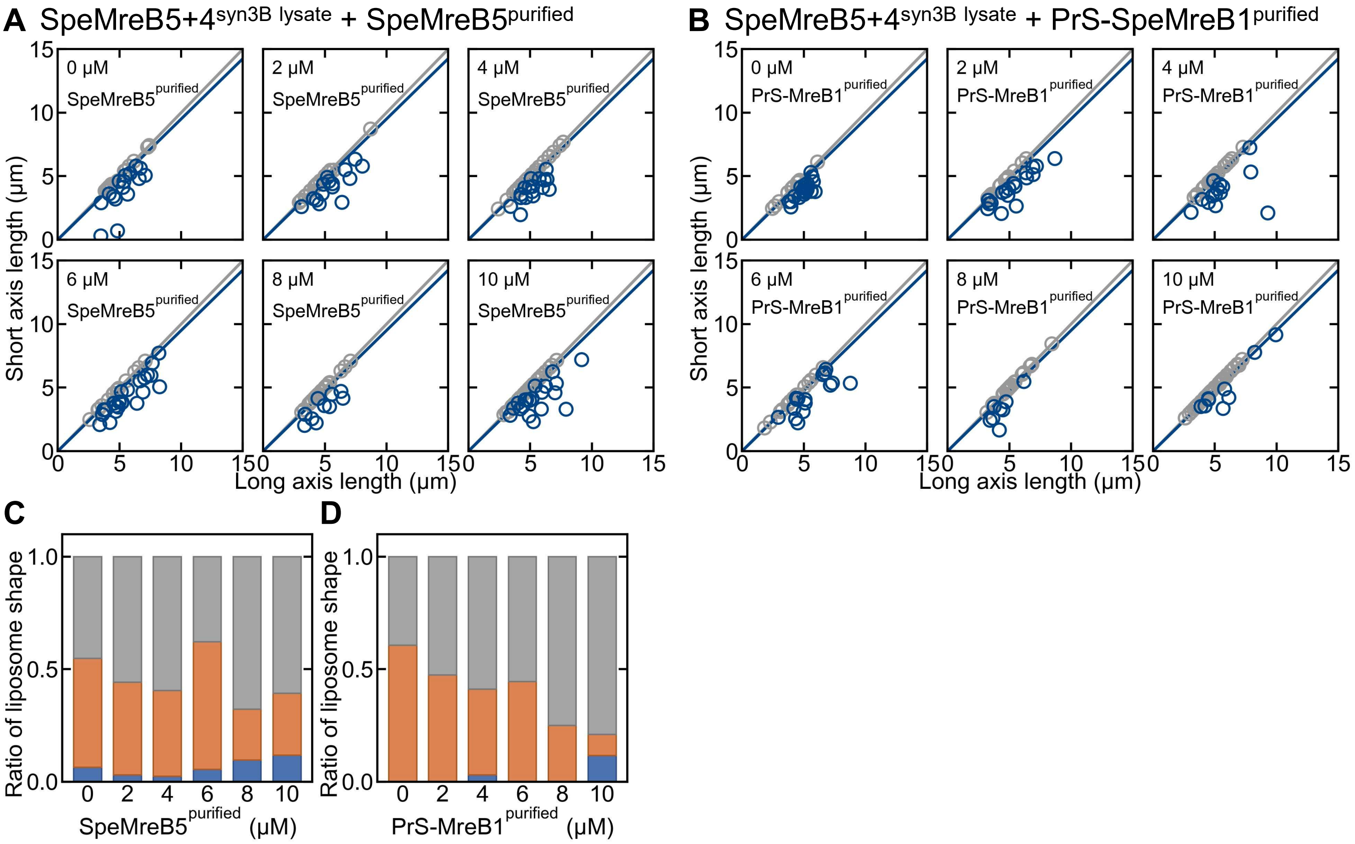

A SpeMreB5+4syn3B lysate + SpeMreB5purified
B SpeMreB5+4syn3B lysate + PrS-SpeMreB1purified
0 μM SpeMreB5purified
2 μM SpeMreB5purified
4 μM SpeMreB5purified
6 μM SpeMreB5purified
8 μM SpeMreB5purified
10 μM SpeMreB5purified
0 μM PrS-MreB1purified
2 μM PrS-MreB1purified
4 μM PrS-MreB1purified
6 μM PrS-MreB1purified
8 μM PrS-MreB1purified
10 μM PrS-MreB1purified
Short axis length (μm)
Long axis length (μm)
0
5
10
15
C
D
Ratio of liposome shape
0.0
0.5
1.0
0 2 4 6 8 10
SpeMreB5purified (μM)
PrS-MreB1purified (μM)

### Fig. S5

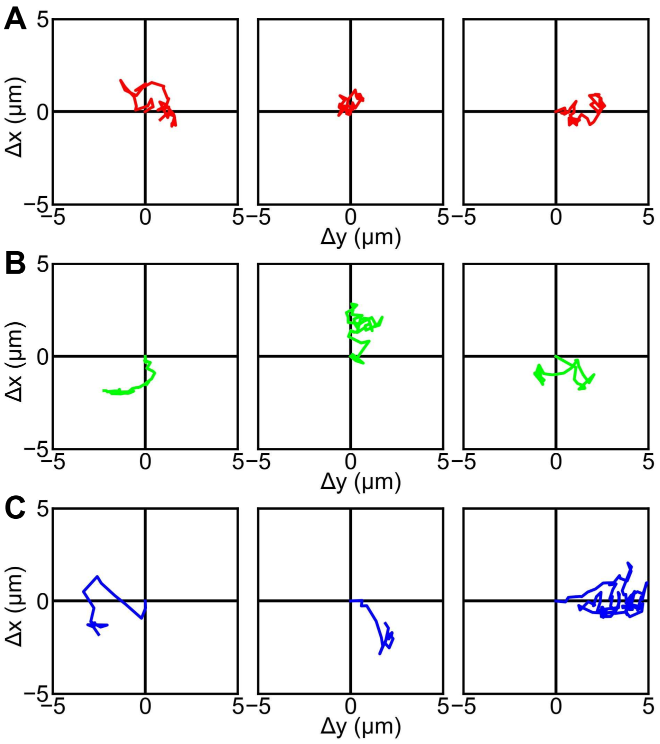
